# Modelling spatial variation in agricultural field trials with INLA

**DOI:** 10.1101/612036

**Authors:** Maria Lie Selle, Ingelin Steinsland, John M. Hickey, Gregor Gorjanc

## Abstract

The objective of this paper was to fit different established spatial models for analysing agricultural field trials using the open-source R package INLA. Spatial variation is common in field trials and accounting for it increases the accuracy of estimated genetic effects. However, this is still hindered by the lack of available software implementations. Here we compare some established spatial models and show possibilities for flexible modelling with respect to field trial design and joint modelling over multiple years and locations. We use a Bayesian framework and for statistical inference the Integrated Nested Laplace Approximations (INLA) implemented in the R package INLA. The spatial models we use are the well-known independent row and column effects, separable first-order autoregressive (AR1⊗AR1) models and a geostatistical model using the stochastic partial differential equation (SPDE) approach. The SPDE approach models a Gaussian random field, which can accommodate flexible field trial designs and yields interpretable parameters. We test the models in a simulation study imitating a wheat breeding program with different levels of spatial variation, with and without genome-wide markers, and with combining data over two locations, modelling spatial and genetic effects jointly. We evaluate predictive performance by correlation between true and estimated breeding values, the continuous rank probability score and how often the best individuals rank at the top. The results show best predictive performance with the AR1⊗AR1 and the SPDE. We also present an example of fitting the models to real wheat breeding data and simulated tree breeding data with the Nelder wheel design.

**Key message:** Established spatial models improve the analysis of agricultural field trials with or without genomic data and can be fitted with the open-source R package INLA.

## 1 Introduction

In plant breeding the main goal is to select individuals with the best performance as new market varieties or to select individuals with the best genetic potential as parents of the next generation. To this end breeders use field trials to estimate genetic and breeding values of individuals. Spatial variation is common in such trials and if not accounted for it can impact the estimation. There can be several sources of spatial variation in a field trial, such as changes in fertility, watering and soil depth. Other sources of spatial variation that often occur are external influences due to the way plots are treated, for example the effect of drilling, spraying and harvesting. This extraneous variation can be handled by the addition of further effects in a model, such as column or row effects.

Traditionally, spatial variation has been accounted for by using control plots, replications and blocks. These approaches do not account for all spatial variability, in particular they do not account for dependency between neighbouring blocks and plots within blocks, which can affect the estimation of genetic values. Several models have been proposed to model spatial variation. One of the most widely used is the separable first-order autoregressive (AR1⊗AR1) model introduced by Cullis and Gleeson (1991) and extended by Gilmour et al. (1997). It has been shown to fit well in many trials (e.g., Gilmour et al., 1997; Rodríguez-Álvarez et al., 2018). There are other models that can correct for spatial variation. For example, there is a whole class of Gaussian intrinsic models based on the seminal work of Besag and Higdon (1999), which have not gained much traction in plant breeding applications. Much has also been done on smoothing techniques, among which the recent SpATS approach explores two-dimensional smooth surfaces through the use of tensor product P-splines (Rodríguez-Álvarez et al., 2018). Nearest neighbour models are reviewed by Piepho et al. (2008) and the use of spatial kernels is also common (Elias et al., 2018).

Most of the popular spatial methods in plant breeding use lags between plot locations as a distance, while continuous spatial variation is not commonly addressed. If observations are irregularly spaced, the autoregressive and other models assuming equal spacing are not applicable. However, there are extensions to the autoregressive model, using covariance functions known as the power model and the exponential model (Schabenberger and Gotway, 2017). The kernel methods presented in Elias et al. (2018) also use covariance functions based on Euclidean distance between plots.

In this paper, we model spatial variation in agricultural field trials using different models with publicly available open-source software. We fit the common column and row effects and the separable first-order autoregressive AR1⊗AR1 model (Cullis and Gleeson, 1991; Gilmour et al., 1997). In addition we fit a Gaussian random field to the field trial through the stochastic partial differential equation (SPDE) approach introduced by Lindgren et al. (2011).

For inference we use the Bayesian numerical approximation procedure known as the Integrated Nested Laplace Approximations (INLA) introduced by Rue et al. (2009) with further developments described in Martins et al. (2013). The method is easy to use through the R package INLA where models are fit with the `inla()` function with the same ease as using the base R functions `lm()` or `glm()`. INLA calculates marginal posteriors for all fixed and random effects, parameters and linear combinations of random effects without using sampling-based methods such as Monte Carlo Markov Chain. It is based on numerical approximations and numerical methods for sparse matrices and is much faster than sampling-based methods (Rue and Martino, 2007).

INLA, and especially the SPDE approach, is flexible with respect to the field trial design and to including several years and locations in the analysis. For example, the SPDE approach can be used beyond the standard lattice design, which we demonstrate with the Nelder wheel design used in forestry (Parrott et al., 2012). For a recent review and comprehensive treatment of INLA and SPDE see (Bakka et al., 2018; Krainski et al., 2018).

The objective of this article was to test established spatial models for analysing agricultural field trials using the open-source R package INLA. INLA and the SPDE approach allows us to fit multi-trial data where designs vary between trials and might even not necessarily be regular. With a simulation study we show that the SPDE approach performs equally well as the AR1⊗AR1 model. Our goal was to show that performing inference with INLA and the SPDE approach is flexible, and yields interpretable parameters for the spatial variation in a field trial. Further, using INLA gives efficient Bayesian inference. We also fitted the models on wheat data from Lado et al. (2013) and on a simulated tree breeding data set with the Nelder wheel design.

## 2 Material and methods

In this section we present the data for a simulated wheat breeding program, a real wheat field trial and a simulated tree breeding trial with the Nelder wheel design. We also present the used statistical models, studied cases, how we inferred model parameters and how we evaluated the different models.

### 2.1 Experimental design and data

#### Simulated wheat data

To evaluate and compare the proposed models, we have simulated a wheat breeding program and corresponding field trials using the R package AlphaSimR (Faux et al., 2016; Gaynor et al., 2019). The simulation followed closely our previous work (Gaynor et al., 2017; Gorjanc et al., 2018), where we simulated a wheat-like genome and 30 years of a wheat breeding program with field trials. In summary the breeding program was based on the following stages:

0. Ancestral simulation of a wheat-like genome with 21 chromosomes, each with 1000 single nucleotide polymorphism markers and 1000 quantitative trait loci.
1. 50 parental inbred lines.
2. 100 crosses between the parental lines with 100 doubled-haploid lines per cross, resulting in a total 10,000 lines.
3. Plant the 10,000 doubled-haploid lines in headrows and observe a phenotype that is correlated to yield and had a heritabiliy of 0.03.
4. Select 1,000 best individuals from the headrows and plant them in a preliminary yield trial for the first yield phenotype assessment with 0.25 heritability.
5. Select 100 best individuals from the preliminary yield trial and plant them in an advanced yield trial for the second yield phenotype assessment with 0.45 heritability.
6. Select 10 best individuals from the advanced yield trial and plant them in an elite yield trial for the final yield phenotype assessment with 0.62 heritability to release the best individual as a commercial variety.
7. Select 50 best individuals from preliminary, advanced and elite yield trials as parents of the next generation.

To simulate yield phenotypes we summed the year, location, individual genetic, spatial dependent plot and independent plot residual effects. We have focused our attention to the preliminary yield trial (4.) only, because this stage has low replication, which makes modelling spatial variation important. The 1000 lines in the preliminary yield trial were planted in two locations, each with plots arranged in a lattice with 50 rows and 20 columns. The distance between columns was twice as large as the distance between rows due to long and narrow plot shape. We sampled year and location effects from a Gaussian distribution with an expected value of 0 and variance equal to residual variance. Individual genetic effects were based on quantitative trait loci genotypes and corresponding allele substitution effects (Faux et al., 2016; Gaynor et al., 2019). We sampled plot spatial effects from a Gaussian random field generated from an SPDE model with a spatial range of 10 units. We varied the proportion of variation due to spatial effects to be 0%, 50%, or 75% of the residual variance, that is, with 50% a half of variation between plots was due to spatial effects and a half due to other unknown effects (plot residual). More detailed description of the Gaussian random field and the SPDE approach is given in Section 2.2.2 and 2.4.1. We standardized the data before data analysis.

#### Chilean wheat data

We used parts of the wheat field trial data presented in Lado et al. (2013) and used by Rodríguez-Álvarez et al. (2018) as shown in Fig. 1. The data consisted of 384 advanced lines from wheat breeding programs in Chile and Uruguay in years 2011 and 2012. We analysed the total grain yield harvested within each plot. Plots were twice as long as wide. The experimental design was an alpha design with two replicates. The trial had 40 rows and 20 columns and according to Rodríguez-Álvarez et al. (2018) the replicates were placed such that the first/second 20 rows corresponded to the first/second replicate. This is indicated by the horizontal line in Fig. 1.

**Fig. 1.**
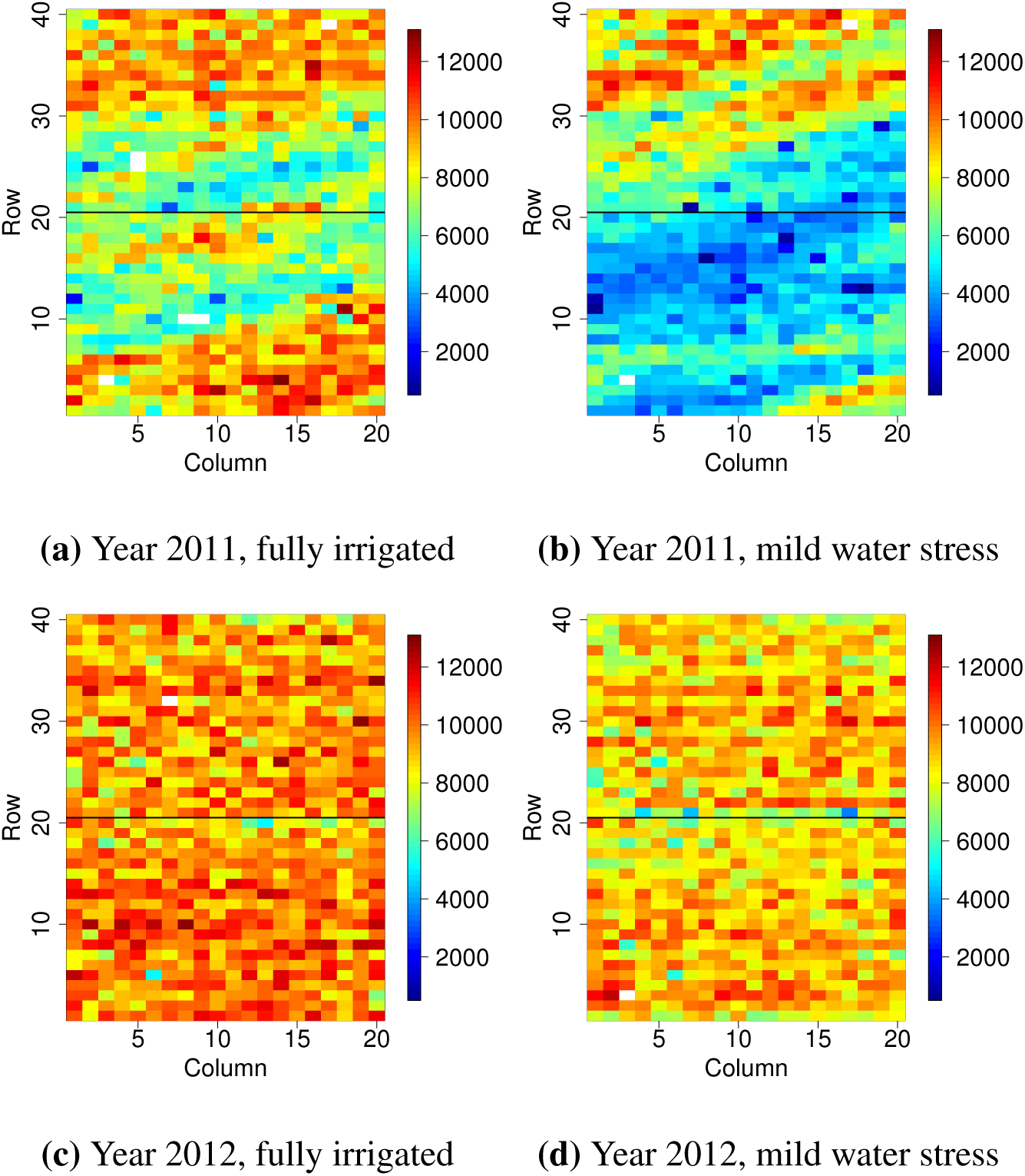
Grain yield in the Chilean wheat data (Lado et al., 2013)

The advanced lines were evaluated in the Santa Rosa region under two different levels of water supply: full irrigation and mild water stress. This gave four data sets each with 800 observations. The 384 lines had also 102,324 genome-wide markers. We imputed missing genotypes with the average allele dosage and computed the VanRaden (2008) genomic relationship matrix among the 384 advanced lines. In addition to the 384 advanced lines, phenotype observations were available for additional 16 lines that were not genotyped. We assumed that these had a genomic relationship of zero between themselves and the 384 advanced lines. We standardized the data before data analysis.

#### Simulated tree data with the Nelder wheel design

We also simulated data with a design used by tree breeders to test the effect of multiple planting densities on tree growth, known as the Nelder wheel design (Parrott et al., 2012). We chose this particular design to show the flexibility of the R package INLA and the SPDE approach. The Nelder wheel design is circular with rings radiating outward with increasing distance. Spokes connect the center with the furthest ring, and at the intersections of spokes and rings, a tree is planted. The variable planting densities within a single trial eliminates the need for separate trials for each planting density.

In the simulation we tested 10 different planting densities with 30 planted trees for each density. The inner circle had a radius of 10, and the 9 subsequent circles had a radius of 1.15 times the radius of the previous circle (Fig. 2).

**Fig. 2.**
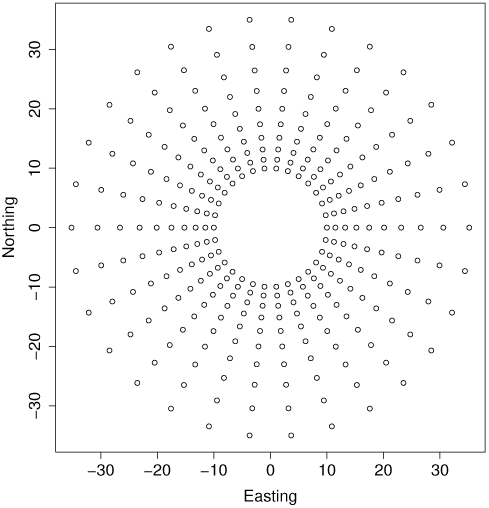
Depiction of the Nelder wheel plot design

We simulated the phenotype for each tree as a sum of the intercept with a value of 10, the tree density covariate multiplied by a regression coefficient of 10, a spatial effect simulated from a Gaussian random field using the SPDE approach, and a Gaussian residual with zero mean and variance 0.5. The simulated field (spatial effects) had a variance of 0.5 and a range of 10.

The growing area available to each tree *i* was calculated from:

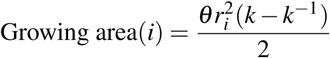

where *θ* was the angle between arcs in radians and *r*_*i*_ was the radius of circle *i*. The factor *k* was 1.15. The planting density was then calculated as the inverse of the growing area.

### 2.2 Statistical models

We assumed to have *n* plots such that a single field trial was indexed by the rows and columns of an *r × c* array. There were *m ≤ n* different genetic lines planted in these plots. The observed phenotype *y*(*s*_*i*_) was assumed to be a realization of a random variable *Y* (*s*_*i*_) in plot coordinates *s*_*i*_ ∈ ℝ^2^, *i* = 1, …, *n*. We considered the following general additive linear model:

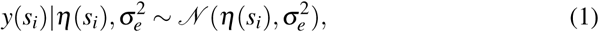

with

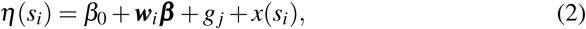

where *β*_0_ was an intercept, 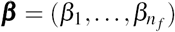 was a vector of effects with a known covariate vector ***w***_*i*_ for plot *i* with 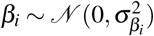, *g* _*j*_ was the genetic effect for individual *j* = 1, *…, m* tested in the plot *i* and *x*(*s*_*i*_) was the spatial effect for the plot.

#### 2.2.1 Genetic effect

We assumed that the genetic effect *g* _*j*_ was a sum of an additive genetic effect (breeding value) *a* _*j*_ and a non-additive (residual) genetic effect *n* _*j*_. For non-additive genetic effects we assumed an independent prior distribution 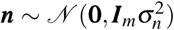. For additive genetic effects we assumed that they were fully explained by genome-wide markers such that 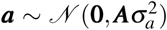, where ***A*** is a relationship matrix. We calculated the relationship matrix as ***A*** = ***ZZ***^*T*^ */k*, where ***Z*** was a column-centred genotype matrix of dimension *m × p, p* was the number of markers, and *k* = 2 ∑_*l*_ *q*_*l*_ (1 *-q*_*l*_) with *q*_*l*_ being allele frequency at marker *l* (VanRaden, 2008). An equivalent model for the additive genetic effects was to use the genotype matrix directly, letting ***a*** = ***Zu***, where ***u*** were marker effects 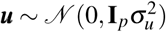.

Genome-wide marker data contains substantial amount of shared information among related individuals due to shared genome segments. Therefore, we could compress it to reduce model dimension while retaining information, which saved computation time (e.g., Jolliffe, 1986). With singular value decomposition we obtained:

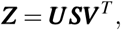

where ***U*** was a unitary matrix of dimension (*m× m*), ***S*** was the diagonal matrix (*m× n*) of singular values and ***V*** was an (*n × n*) matrix of eigenvectors. We used the principal components (the columns of *Z****V***) corresponding to the largest singular values of ***S*** and chose *p\**components that explained approximately 95% of the variation in ***Z***. That is, we replaced the ***Z*** by ***Z****\**= *Z****V*** (:, 1 : *p\**) of dimension *m × p\**. The linear predictor from (2) then became:

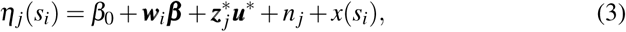

where 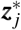 was the *j*-th row vector of ***Z****\**for individual *j* and 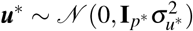 were principal component effects.

#### 2.2.2 Spatial effect

We tested the independent row and column effects model, the separable first-order autoregressive (AR1⊗AR1) model, and a Gaussian random Markov field through the SPDE approach. The independent row and column model and separable autoregressive model are based on a discretisation of the field and model only a finite collection of spatial random variables. For these models we omit the *s*_*i*_ in *x*(*s*_*i*_) and use *x*_*i*_. This is to emphasize that these models use neighbouring plots as opposed to the SPDE approach which is a continuous spatial process and for which we use the notation *x*(*s*_*i*_).

##### Row and column effects model

Row and column effects can model the underlying smooth spatial field as well as external variation due to field management. We assumed:

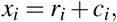

where 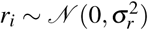 was the row effect and 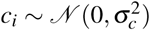 was the column effect of plot *i, i* = 1, *…, n*.

##### Separable autoregressive model, AR1⊗AR1

The autoregressive model of order 1 (AR1) for the Gaussian vector ***x*** = (*x*_1_, *…, x*_*r*_) is defined as:

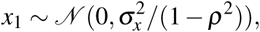

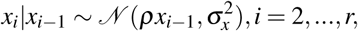

where |*ρ*| < 1.

For modelling the influence of neighbouring plots along rows and columns, the autoregressive model in each direction was combined into a two-dimensional first-order separable autoregressive model (Cullis and Gleeson, 1991; Gilmour et al., 1997), denoted as AR1⊗AR1. In this model the spatial effect vector ***x*** of length *n* was modelled as:

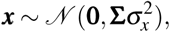

with **Σ** = **Σ**_*r*_ ⊗ **Σ**_*c*_. The matrices **Σ**_*r*_ and **Σ**_*c*_ were the covariance matrices of first-order autoregressive processes in row and column direction respectively, and ⊗ was the Kronecker product. The model had two dependency parameters, one in each direction, *ρ*_*r*_ and *ρ*_*c*_, and a variance parameter 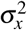.

##### Gaussian random fields

In the model described above, the spatial variation was modelled as discrete. Assuming a continuous field for the spatial variation was however more realistic. Continuously indexed Gaussian random fields play an important role in spatial statistical modelling and geostatistics. In the field *𝒟* ∈ ℝ^*d*^ with coordinates ***s*** ∈ *𝒟*, the continuously indexed Gaussian random field *x* (***s***) has a joint Gaussian distribution for all finite collections *{x*(***s***_*i*_)*}*. The Gaussian random field is specified through the mean ***µ*** and the covariance matrix **Σ** = *C*(***s***_*i*_, ***s*** _*j*_).

In this study we used the Matérn covariance function, which is the most important covariance function in spatial statistics (Stein, 2012). The Matérn covariance function between locations ***s***_*i*_, ***s*** _*j*_ ∈ ℝ^*d*^ was:

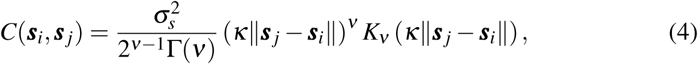

where *K*_*v*_ was the modified Bessel function of the second kind and order *v >* 0. The parameter *κ* can be expressed as 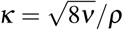, where *ρ >* 0 was the range parameter describing the distance where the correlation between two points was near 0.1, and 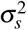 was the marginal variance. The parameter *v* determined the mean-square differentiability of the field. The SPDE approach is a computationally efficient way to fit a Gaussian random field (Lindgren et al., 2011), which we describe in 2.4.1.

#### 2.2.3 Prior distributions

We used a full Bayesian approach to estimation which requires prior distributions for all parameters. We have already specified prior distribution for most location (mean) parameters. We used the default priors of INLA, that is, a Gaussian prior with mean 0 and variance 1000 for the intercept and covariate effects and an inverse gamma prior with shape 1 and inverse scale 5 *×* 10^*-*5^ for variance parameters. In the separable autoregressive model the inverse gamma prior was set for the marginal variance 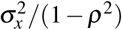, and for *ρ*, the transformed variable log((*ρ* + 1)(*ρ -* 1)) was assigned a Gaussian prior with mean 0 and standard deviation 0.15. For the SPDE approach priors were specified for the parameters *κ* and *τ* that control spatial range and variance, see 2.4.1. We used the default joint Gaussian prior on log(*κ*) and log(*τ*) with mean 0 and identity covariance matrix, so that log(*κ*) and log(*τ*) were independent (Blangiardo and Cameletti, 2015).

### 2.3 Case studies

#### 2.3.1 Simulation study

We fitted the model (1) with two versions of the linear predictor (3) to the preliminary yield trial of each simulated breeding program - without and with genome-wide markers. The two linear predictors were:

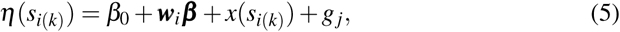

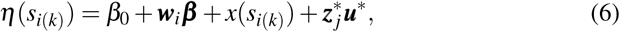

where 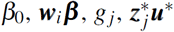 were as described as in the Section 2.2. The linear predictors differed in that model (5) assumed individuals were genetically independent, whereas model (6) used genome-wide marker data to model the genetic dependency. The linear predictors included both trials simultaneously. The subscript *k* = 1, 2 in *s*_*i*(*k*)_ indicated that the plot coordinates *s*_*i*_ were in field *k* and a fixed effect of a field was included in ***w***_*i*_ ***β***. Otherwise the *k* fields were assumed to be independent realizations from the same distribution, and we used all three spatial models described in Section 2.2.2 to fit spatial variation. We also fitted a model where the spatial effect was omitted, which we denoted as the NoSpatial model. Since distance between columns were twice as large as the distance between rows, we accounted for this in the SPDE approach, by appropriate scaling the column coordinates. The matrix *Z\**was constructed using *p\**= 500 principal components of the singular value decomposition of the centered genotype matrix *Z*.

#### 2.3.2 Chilean wheat data

Using the four data sets from Lado et al. (2013) presented in Section 2.1, we fitted the model (1) with different versions of the linear predictor (3). The four linear predictors were:

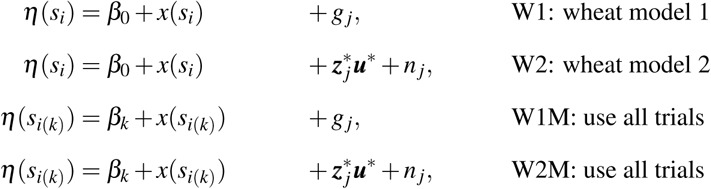

where 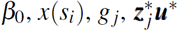, and *n* _*j*_ were as described in Section 2.2. As with the simulation study we used the three spatial models described above and the NoSpatial model. The linear predictors W1M and W2M included all four trials simultaneously and therefore the intercept *β*_*k*_, *k* = 1, *…,* 4, was trial specific. Further, the subscript *k* in *s*_*i*(*k*)_ indicated that the plot coordinates *s*_*i*_ were in field *k*. The four trials in Fig. 1 showed quite different spatial patterns, so it was not reasonable to assume that they were realizations from the same distribution. We therefore modelled the spatial effect in the trials from 2011 as independent realizations from the same underlying distribution, and the same for the 2012 trials. This gave two sets of spatial parameters in the model, one set for the 2011 trials and one set for the 2012 trials. The matrix *Z\**was constructed using *p\**= 280 principal components of the singular value decomposition of the centered and scaled genotype matrix *Z*.

#### 2.3.3 Nelder wheel plot

To analyse the simulated tree data, we fitted the model (1) with the following linear predictor:

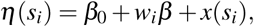

where *β*_0_ was the intercept, *β* was a density effect and the spatial effect *x*(*s*_*i*_) was modelled as a Gaussian random field using the SPDE approach.

### 2.4 SPDE, Inference and Evaluation of case studies

#### 2.4.1 The SPDE approach to spatial modelling

Modelling with Gaussian random fields is computationally challenging because they give rise to dense precision matrices that are numerically expensive to factorise in the estimation procedures (Rue and Held, 2005). Gaussian Markov random fields do not incur this penalty because they have a sparse precision matrix due to their Markov property. Lindgren et al. (2011) showed how to construct an explicit link between (some) Gaussian random fields and Gaussian Markov random fields by showing that the approximate weak solution of the SPDE:

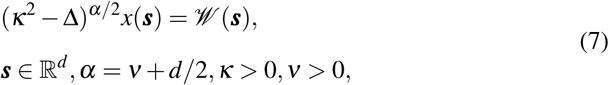

is a Gaussian random field with Matérn covariance function as given in (4). Here, *𝒲* (*·*) is Gaussian white noise, Δ is the Laplacian, *α* is a smoothness parameter, *κ* is the scale parameter in (4), *d* is the dimension of the spatial domain and *τ* is a parameter controlling the variance. The parameters of Matérn covariance are linked to the SPDE through:

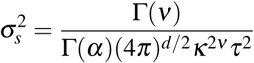

where *v* = *α* −*d*/2, and we use *α* = 2 and *d* = 2.

A Gaussian Markov random field approximation described in Lindgren et al. (2011) is enabled by solving the SPDE in (7) by the finite element method. Further details on the SPDE approach to spatial modelling can be found in Lindgren et al. (2011).

#### 2.4.2 Bayesian inference with INLA and the INLA package

Statistical inference is carried out using the INLA methodology introduced in Rue et al. (2009), which is implemented for use in `R` (R Core Team, 2018) in the package `INLA` (available at www.r-inla.org). In this section we give a short introduction to the class of models known as latent Gaussian models and how INLA can be used to approximate the posterior marginal distributions for such models. For an in-depth description of INLA we refer to Rue et al. (2009), Martins et al. (2013) and the recent review by Rue et al. (2017).

Latent Gaussian models are hierarchical models in which observations ***y*** are assumed to be conditionally independent given a latent Gaussian random field ***x*** and hyper-parameters ***θ*** _1_, that is *π*(***y****|****x***, ***θ*** _1_) *∼* Π_*i∈ ℐ*_ *π*(*y*_*i*_*|x*_*i*_, ***θ*** _1_). The latent field ***x*** includes both fixed and random effects and is assumed to be Gaussian distributed given parameters ***θ*** _2_, that is *π*(***x****|****θ*** _2_) *∼ 𝒩* (***µ***(***θ*** _2_), **Σ**(***θ*** _2_)). The parameters ***θ*** = (***θ*** _1_, ***θ*** _2_) are known as hyper-parameters and control the Gaussian field and the likelihood for the data. These parameters must be assigned a prior density to specify the latent Gaussian model completely. The class of latent Gaussian models includes many models, for example generalised linear (mixed) models, generalised additive (mixed) models, and spline smoothing methods.

The main aim of Bayesian inference is to estimate the marginal posterior distribution of the variables in the model, that is *π*(*θ*_*j*_*|****y***) for hyper-parameters and *π*(*x*_*i*_*|****y***) for location parameters. INLA computes approximations to these densities quickly and with high accuracy. Laplace approximations are applied to integrals that are Gaussian or close to Gaussian, and for non-Gaussian problems, conditioning is done to break down the approximations into smaller sub-problems that are almost Gaussian.

For the computations in INLA to be both quick and accurate, the latent Gaussian models have to satisfy some additional assumptions. Since INLA integrates over the hyper-parameter space, the number of non-Gaussian hyper-parameters ***θ*** should be low, typically less than 10, and not exceeding 20. Further, the latent field should not only be Gaussian, it must be a Gaussian Markov random field. The conditional independence property of a Gaussian Markov random field yields sparse precision matrices which makes computations in INLA fast due to efficient algorithms for sparse matrices. Lastly, each observation *y*_*i*_ should depend on the latent Gaussian field through only one component *x*_*i*_.

The INLA package can be downloaded to `R`, and is run using the `inla()` function with three mandatory arguments; a data frame containing the data, a formula much like the formula for the standard `lm()` function in `R`, and a string indicating the likelihood family. The default is Gaussian with identity link. The following call generates an object of type `inla`:

~~~
fit <-inla(data = Data, formula = Formula, family = “Gaussian”)
~~~

Prior distributions are specified through additional arguments. Several tools to manipulate models and likelihoods exist as described in tutorials at the INLA web-page www.r-inla.org and the books by Blangiardo and Cameletti (2015); Krainski et al. (2018). The `R` scripts used for the fitted models and the tree breeding simulation are available in Online Resource 1. Specifically we provide `R` code for all the fitted models to the real wheat data and the simulation and analysis of the tree breeding data with the Nelder wheel design.

Here, we show how to fit an: (i) Row+Col model, (ii) AR1 row and AR1 col model, (iii) AR1⊗AR1 model and (iv) SPDE model. The data should be stored in a data frame or list. Here the data frame `Data` has one row for each observation with columns containing the phenotype, id for each genetic line and row and column in the field. The id for each genetic line is included twice because we want to model the genetic effect with and without genetic markers.

`head(Data)`

| y | IndI | IndMI | Row | Col |
| --- | --- | --- | --- | --- |
| 0.254 | 275 | 275 | 1 | 1 |
| 0.029 | 325 | 325 | 1 | 2 |
| -0.056 | 119 | 119 | 1 | 3 |
| -0.128 | 138 | 138 | 1 | 4 |
| 0.531 | 296 | 296 | 1 | 5 |
| -0.672 | 257 | 257 | 1 | 6 |

In the formula below, we indicate that each genetic line should be modelled both with an independent normal distributed effect, and using marker effects for the markers stored in `Gen`, the approach described in Section 2.2.1.

~~~
Formula <-y ∼ f(IndI, model = “iid”) + f(IndMI, model = “z", Z = Gen)
~~~

To include a spatial model, one of the following functions can be added to `Formula`.

~~~
(i): f(Row, model = “iid”) + f(Col, model = “iid”)
(ii): f(Row, model = “ar1”) + f(Col, model = “ar1”)
(iii): f(Row, model = “ar1", group = Col, control.group = list(model = “ar1”))
(iv): f(Field, model = Spde)
~~~

The models with formula including either of effects (i) - (iii) are fitted with the call to `inla()` as described above. The SPDE model (iv) requires a few additional stages which we show in the full code available in Online Resource 1.

#### 2.4.3 Evaluation of model performance

We evaluated the models using the correlation between the true and estimated values, the continuous rank probability score (CRPS), by identifying the top individuals, and the residual variance.

We used the CRPS to take into account the whole posterior predictive distribution, that is, to compare the estimated posterior means with the true/observed values while accounting for the uncertainty of estimation. The CRPS is defined as (Gneiting and Raftery, 2007):

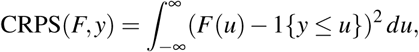

where *F* is the cumulative distribution of the estimator of interest and *y* is the observed value. The CRPS is negative oriented, so the smaller the CRPS the closer the estimated value is to the observed/true value. For readers not familiar with the CRPS, three plots in Fig. 3 show the cumulative distribution functions for estimates and the observed value of 1.0. In Fig. 3a the estimate is close to the true value and the area between the curves is small and so is the CRPS. In Fig. 3b the estimated mean is equal to the true value, but the large uncertainty due to estimation causes a large area between the curves, and hence a larger CRPS than in Fig. 3a. In Fig. 3c the uncertainty of the estimation is small, but the estimated mean is further from the true value, causing the area and the CRPS to be large.

**Fig. 3.**
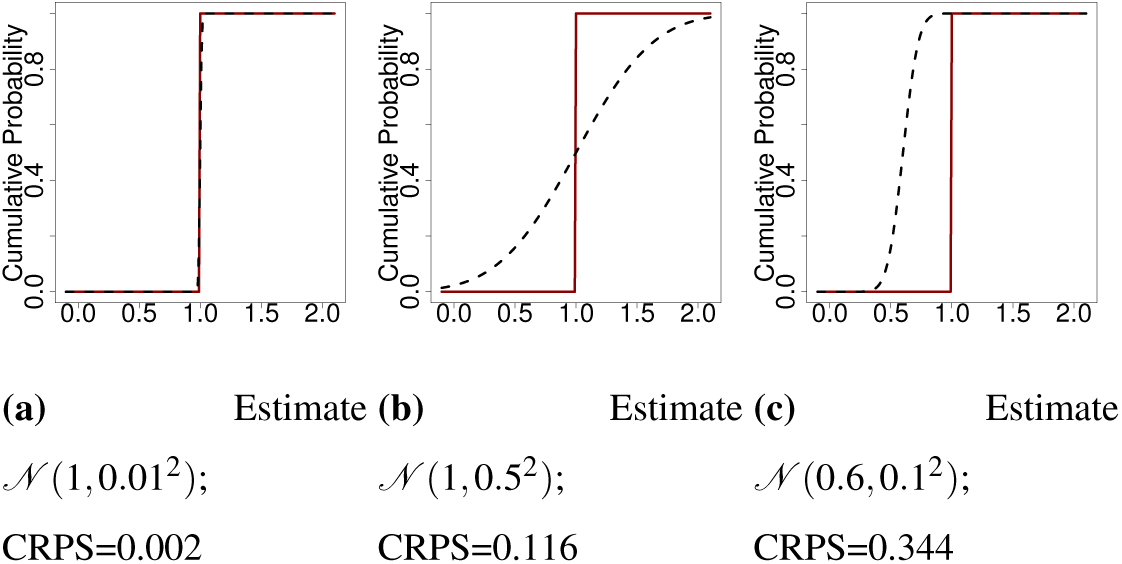
Cumulative distribution function (CDF) of the observation (true value = 1; solid line) and of estimate (dashed line)

For the simulated data we computed the correlation and the CRPS between true and estimated breeding value. We also quantified how many of the ten best individuals were among the estimated top 100 individuals.

For the real data we did not know the true breeding value, and it was therefore not possible to validate the estimated breeding values. We therefore focused on the residual variance from each model as a measure of the unexplained variance. This value can be seen as a proxy for the coefficient of determination (*R*^2^), a measure on how much of the data variance is explained by a given model (Gelman and Hill, 2006).

## 3. Results

In this section we present the results from the three cases presented in the Section 2.3. In the results from the simulation study we compare correlation, CRPS and top ranking of individuals between the spatial models. In the results from the real data we present estimated genetic variances, marker variances and residual variances and compare these between the different models. In the results from the simulated tree breeding data we present the posterior distribution of all parameters and the estimated spatial effect.

### 3.1 Simulation study

This section presents the results from the simulation study. The models were evaluated using the correlation and CRPS between the true and estimated breeding value and using the number of the top ten individuals that were among the top 100 ranked individuals when considering estimated breeding value (posterior mean). In this section all tables have three scenarios indicating the proportion of environmental variance due to spatially structured variation in the data, 0.00, 0.50 or 0.75, while the total variance was the same. Proportion of spatial variation therefore indicates how much of total environmental variance was due to structured spatial noise and unstructured noise (see Section 2.1). Tables showing average estimates for residual variance, genetic and marker variance and other spatial hyper-parameters are given in the Online Resource 2.

In Table 1 the average correlation is presented, in Table 2 the average CRPS is presented and in Table 3 the average number of the top ten individuals that are among the top 100 ranked individuals is presented. The average was taken over 100 independent replications of the breeding program described in the Section 2.1. We note that genomic data improved the correlation, CRPS and the average number of the top ten individuals for all models and proportions of spatial variance. We further note that modelling the spatial variation also improved these metrics. Below we go though each table in detail.

**Table 1:**
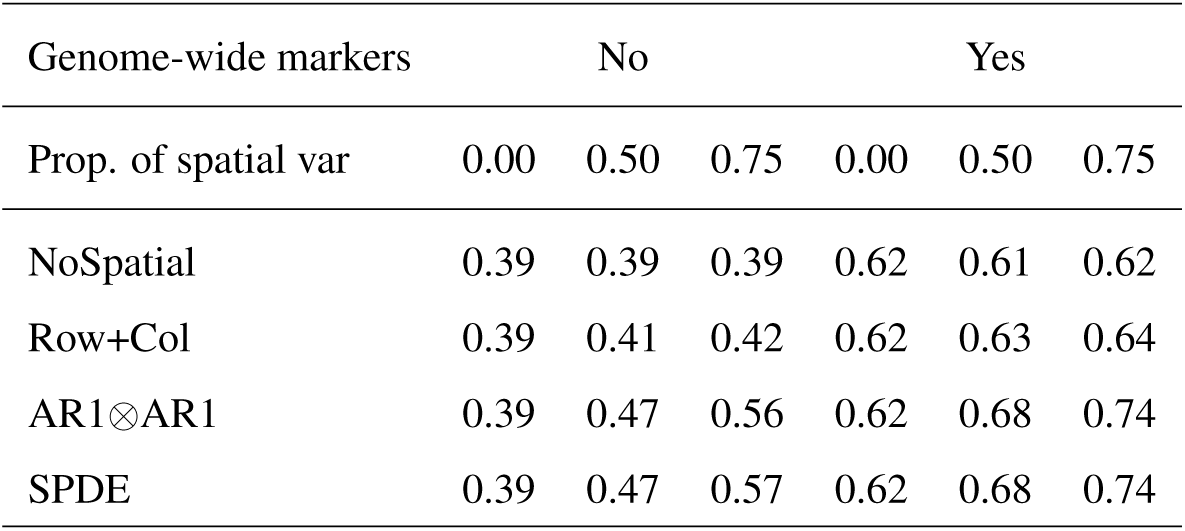
Correlation between the true and estimated breeding value in the preliminary yield trial by the proportion of spatial variation, the spatial model and using genome-wide markers. The standard error was around 0.002

**Table 2:** CRPS between the true and estimated breeding value in the preliminary yield trial by the proportion of spatial variation, the spatial model and using genome-wide markers. The standard error was around 0.0002

| Genome-wide markers | No |  |  | Yes |  |  |
| --- | --- | --- | --- | --- | --- | --- |
| Prop. of spatial var | 0.00 | 0.50 | 0.75 | 0.00 | 0.50 | 0.75 |
| NoSpatial | 0.149 | 0.149 | 0.149 | 0.114 | 0.115 | 0.114 |
| Row+Col | 0.169 | 0.142 | 0.138 | 0.114 | 0.113 | 0.111 |
| AR1$\otimes$AR1 | 0.169 | 0.127 | 0.117 | 0.114 | 0.108 | 0.100 |
| SPDE | 0.148 | 0.127 | 0.117 | 0.114 | 0.107 | 0.099 |

**Table 3:** Average number of the top ten individuals among the top 100 ranked individuals in the preliminary yield trial by the proportion of spatial variation, the spatial model and using genome-wide markers. The standard error was around 0.05

| Genome-wide markers | No |  |  | Yes |  |  |
| --- | --- | --- | --- | --- | --- | --- |
| Prop. of spatial var | 0.00 | 0.50 | 0.75 | 0.00 | 0.50 | 0.75 |
| NoSpatial | 3.89 | 3.82 | 3.89 | 6.32 | 6.41 | 6.43 |
| Row+Col | 3.89 | 4.05 | 4.20 | 6.33 | 6.59 | 6.77 |
| AR1$\otimes$AR1 | 3.89 | 4.81 | 5.81 | 6.32 | 7.36 | 8.07 |
| SPDE | 3.89 | 4.80 | 5.85 | 6.32 | 7.38 | 8.15 |

Across all metrics the SPDE and AR1⊗AR1 stand out as the best models to model the spatial variation. These had the highest correlation when spatial variation was present as seen in Table 1. When there was no spatial variation, the two models did not perform worse than not including a spatial effect. The performance increased as the extent of spatial variation increased. The CRPS results in Table 2 show lower CRPS for the SPDE and the AR1⊗AR1 models compared to the NoSpatial and Row+Col models. These results are in line with the correlation results with one exception for the AR1⊗AR1 model. We also note an improvement in CRPS with increasing extent of spatial variation.

The average times the ten best individuals were among the top 100 ranked individuals are given in Table 3. The SPDE and AR1⊗AR1 models again had better results when there was spatial variation in the data and when genome-wide markers were used in this setting there were on average between 6 and 8 of the top 10 individuals among the top 100 ranked individuals. As expected, the NoSpatial showed no improvement when the degree of spatial variation was increased and the Row+Col model showed only a little improvement with respect to all evaluations.

We also evaluated predictions of breeding values for 1000 doubled-haploid individuals that were genotyped, but not phenotyped. These individuals served to test out-of-sample prediction, which we could perform using estimated genome-wide marker effects. The average correlation between the true and predicted breeding value is presented in Table 4, where AR1⊗AR1 and SPDE again had the highest correlation. For the CRPS in Table 5 we see a similar trend as for the phenotyped individuals, however the improvement with the higher degree of spatial variation is now less dominant. Finally, the average number of the top ten individuals among the 100 ranked individuals is given in Table 6. These results improved with the SPDE and AR1⊗AR1 models and with the increasing spatial variation. The results for the non-phenotyped doubled-haploid lines showed lower correlation, higher CRPS and lower number of the top ten individuals captured than in the preliminary yield trial. This is expected as we had not observed any phenotype data on the doubled-haploid lines.

**Table 4:**
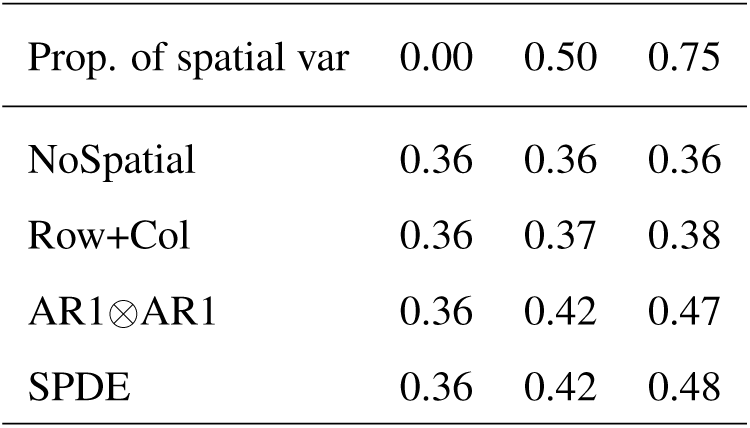
Correlation between the true and predicted breeding value for the non-phenotyped doubled-haploid lines by the proportion of spatial variation and the spatial model. The standard error was around 0.004

**Table 5:** CRPS between the true and predicted breeding value for the non-phenotyped doubled-haploid lines by the proportion of spatial variation and the spatial model. The standard error was around 0.00004

| Prop. of spatial var | 0.00 | 0.50 | 0.75 |
| --- | --- | --- | --- |
| NoSpatial | 0.128 | 0.128 | 0.129 |
| Row+Col | 0.128 | 0.128 | 0.127 |
| AR1$\otimes$AR1 | 0.128 | 0.126 | 0.122 |
| SPDE | 0.128 | 0.126 | 0.122 |

**Table 6:** Average number of the top ten individuals among the top 100 ranked individuals for the non-phenotyped doubled-haploid lines by the proportion of spatial variation and the spatial model. The standard error was around 0.06

| Prop. of spatial var | 0.00 | 0.50 | 0.75 |
| --- | --- | --- | --- |
| NoSpatial | 3.37 | 3.38 | 3.35 |
| Row+Col | 3.37 | 3.51 | 3.60 |
| AR1$\otimes$AR1 | 3.37 | 3.99 | 4.67 |
| SPDE | 3.37 | 3.97 | 4.75 |

### 3.2 Chilean wheat data

In this section we present results from fitting the models W1, W2, W1M and W2M to the Chilean wheat data. We present the estimated genetic variances, marker variances and residual variances from the different spatial models. These are shown in Fig. 4. We also present the posterior predicted phenotype from model W2 for the 2011 trial with full irrigation. All hyper-parameter estimates for the spatial models are given in the Online Resource 2, together with the parameters presented in this section.

**Fig. 4.**
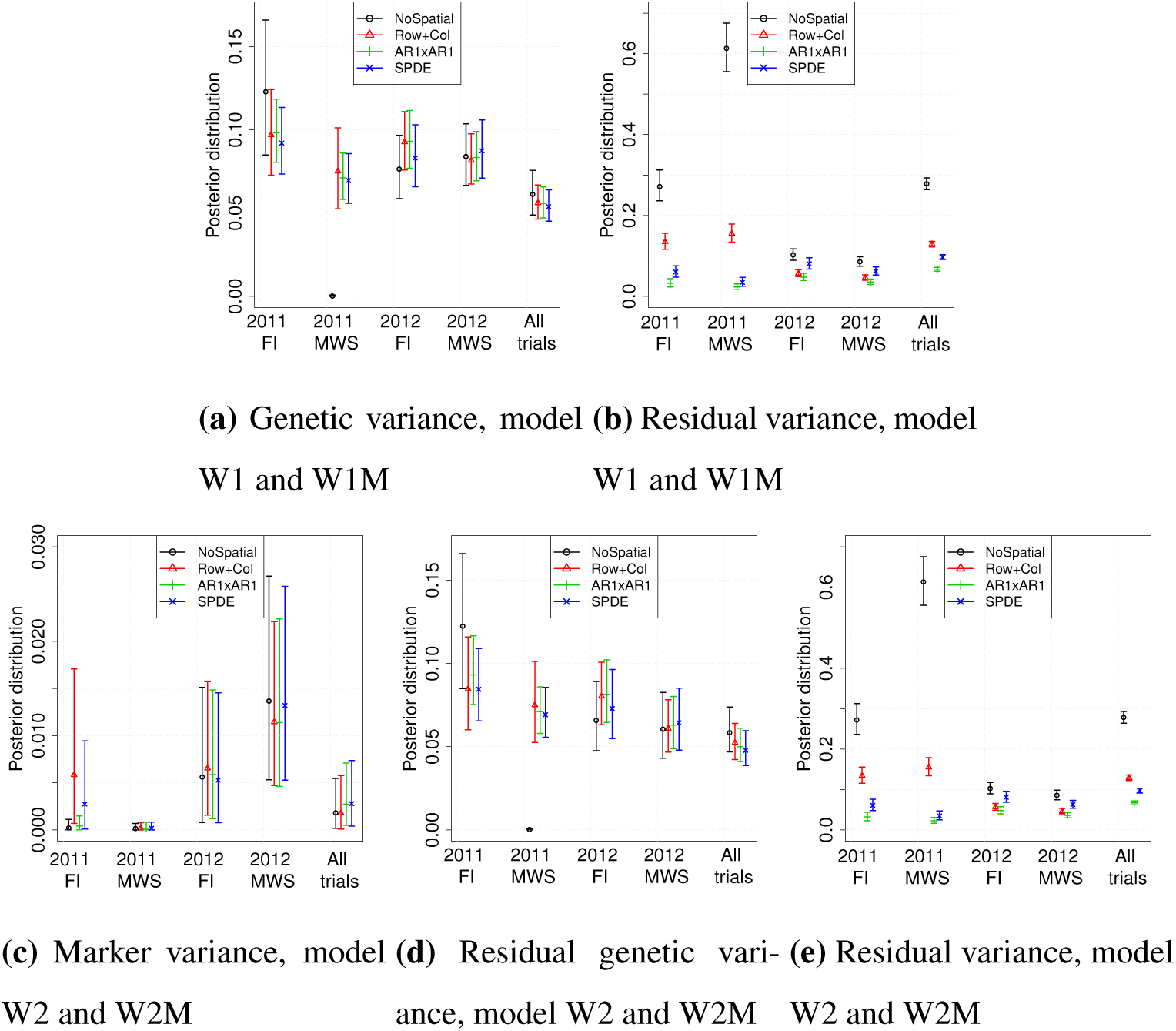
Posterior variances in the models for Chilean wheat data. The top panels are for models that do not use genome-wide marker data (W1 and W1M) and the bottom panels for models that use genome-wide marker data (W2 and W2M)

We first focus on the results from fitting the models without genome-wide markers (models W1 and W1M), which are shown in Fig. 4a and Fig. 4b. The estimated genetic variances were similar within each trial except for the NoSpatial case which assigned all variation to the residual variance in trial 2011 MWS indicating a very bad model fit. Between the trials, there was more variation between the estimates of genetic variance, however most 95% confidence intervals overlap between the different models and trials with a few exceptions. The uncertainty in the genetic variance reduced when all trials were analysed together (W1M), which was expected as more data was used in this model. For the residual variance we expected that it would differ both between models and trials as they described the amount of variation not explained by the structured model terms. As expected the residual variance from NoSpatial was largest as this model cannot explain spatial variation. The AR1⊗AR1 model had the lowest residual variance, closely followed by the SPDE approach in the 2011 trials. When all trials were analysed jointly, the residual variance increased slightly for the AR1⊗AR1 and SPDE approach.

We now focus on the results for models including genome-wide markers (models W2 and W2M) in Fig.4c, Fig. 4d and Fig. 4e. We note that marker variance estimate had large uncertainty and was lower in 2011, particularly in the medium-water stress condition. The genetic variance not captured by markers (Fig. 4d) became more similar between the different trials compared to in Fig. 4a. The residual variance did not change significantly indicating that the markers captured variation that was already captured by the genetic effect modelled in W1 and W1M. However, with genome-wide markers we captured the genetic dependency between individuals with the model, which makes it possible to predict genetic value for non-phenotyped individuals as shown in the previous subsection.

We show the fitted values from model W2 for the 2011 full irrigation trial in Fig. 5. These show how the AR1⊗AR1 model and the SPDE approach managed to capture the spatial pattern in the observations whereas the NoSpatial model and Row+Col model could not. Note that the scale here is different from the one in Fig. 1 since models were fitted to standardized data.

**Fig. 5.**
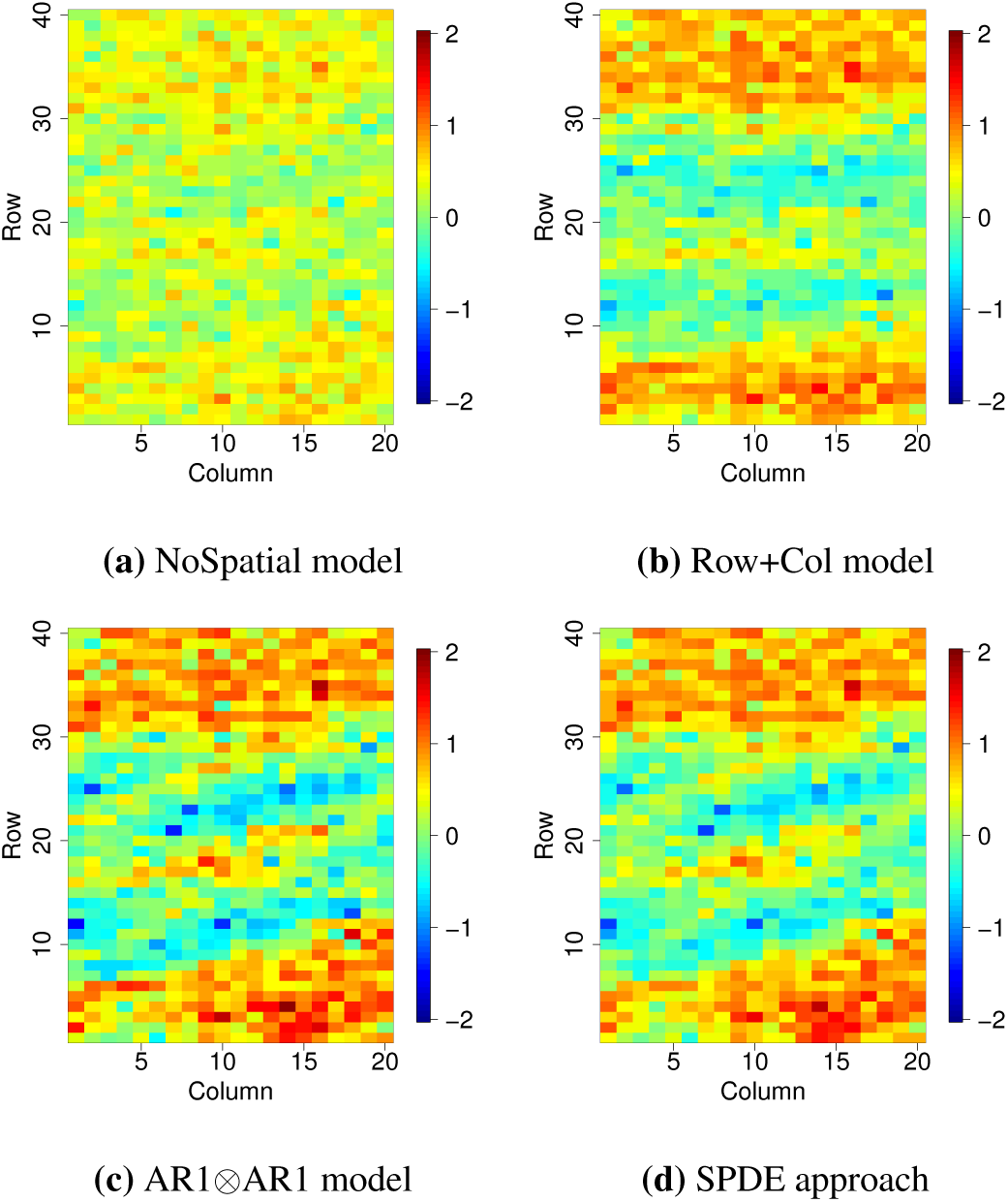
Posterior fitted values from the model W2 for trial 2011 FI using all three methods of spatial correction and no spatial correction.

### 3.3 Nelder wheel plot

In this section, we present the results from fitting the model presented in the Section 2.3.3 to the simulated tree breeding data. In Fig. 6 the posterior distributions for the intercept, fixed density effect, spatial range, spatial variance and residual variance are presented along with the true values used in simulating the data. For all parameters, the posterior distribution contained the true values and the distribution modes were close to the true values.

**Fig. 6.**
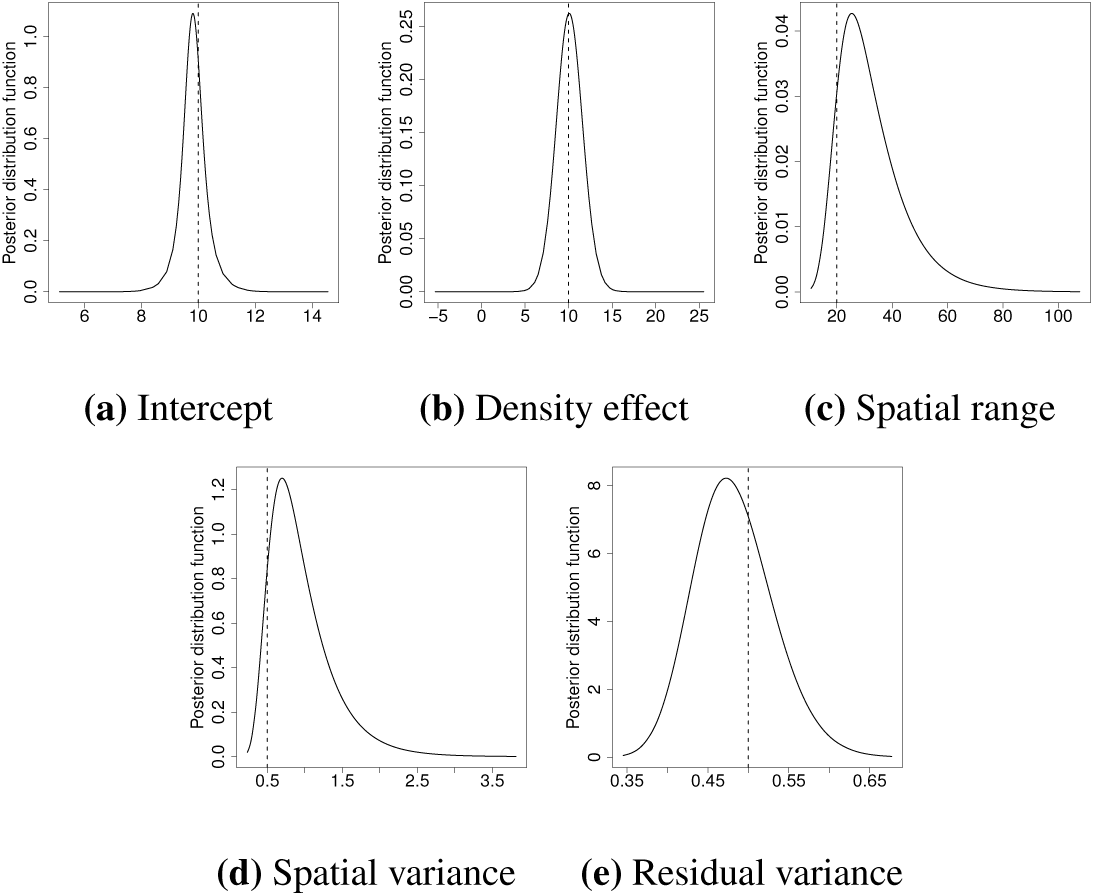
Posterior distributions from model fitted to simulated tree breeding data. Curves represent the posterior distribution and the straight dashed line the true values

In Fig. 7 we show the simulated spatial effect, the posterior mean spatial effect and the standard deviation of the estimate. The mean estimate resembled closely the true spatial field, especially in locations where we had observations. The standard deviation was smallest where we had observations and where the observations were more densely observed.

**Fig. 7.**
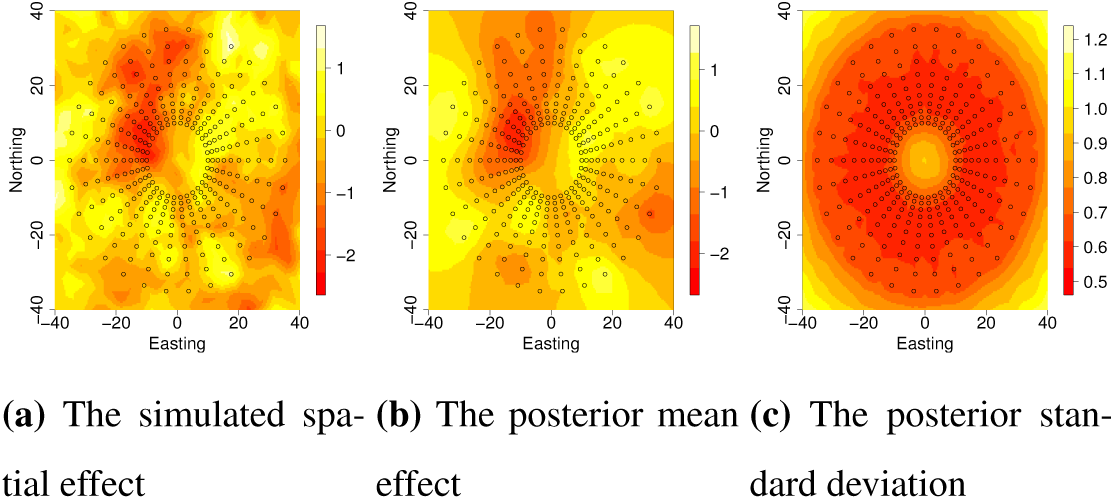
Simulated spatial effect, posterior mean spatial effect and posterior standard deviation of spatial effect in the Nelder wheel example. Black circles indicate tree positions

## 4. Discussion

The objective of this paper was to test established spatial models for analysing agricultural field trials using the open-source R package INLA. We have fitted both spatial and genetic effects jointly in a simulated wheat trial data, a real wheat data set and a simulated tree breeding data set with the Nelder wheel design. Here we highlight three points for discussion: (i) the importance of fitting spatial variation in agricultural field trials, (ii) the flexibility of the R package INLA and the SPDE approach to model multiple trials and years as well as non-standard designs and non-standard phenotype distributions and (iii) the limitations of the R package INLA to estimate large numbers of hyper-parameters, to fit genomic models and with respect to prior specification.

Through the analysis of simulated wheat data sets we showed that the estimates of genetic effects can be improved by accounting for spatial dependency in trials irrespective of the magnitude of the spatial variation. This is in line with the other studies (Elias et al., 2018; Rodríguez-Álvarez et al., 2018; Velazco et al., 2017; Piepho et al., 2008). We observed the greatest improvements with both the AR1⊗AR1 model (Cullis and Gleeson, 1991; Gilmour et al., 1997) and the SPDE approach (Lindgren et al., 2011) as measured using the correlation and continuous rank probability score (CRPS) between the true and estimated effects as well as the average number the top ten individuals that were among the 100 ranked individuals based on the estimates. When we attempted to model non-existing spatial variation, the results were not significantly worse compared to not modelling the spatial dependency. This observation suggests that the AR1⊗AR1 model and the SPDE approach are good default spatial models that do not overfit the data.

Through modelling the real wheat data we demonstrated the flexibility of the R package INLA to model both the genetic and spatial effects for several trials simultaneously. We treated the spatial variation in each trial as an independent realization of the chosen spatial model. By modelling spatial and genetic effects across several trials jointly with one model we did not lose any information as we would if spatial effects were estimated first and then subtracted from the data (Schulz-Streeck et al., 2013). Furthermore, there is a large potential in modelling all trials jointly because this approach enables reduction of the required number of replicates per individual per trial or even individuals per trial(Bernal-Vasquez et al., 2014). It also makes it possible to estimate location and year effects, which can be helpful for future management of the testing sites.

Through modelling the simulated tree data with the Nelder wheel design we demonstrated the flexibility of the SPDE approach with respect to the field trial design. This flexibility arises from the continuous modelling of spatial effects with the SPDE approach as compared to the discrete approach of other standard models. This flexibility is not required for most agricultural field trials, which have a regular lattice layout of plots. The SPDE approach can be used for the regular as well as non-regular designs, which can be useful in special settings, for example, when plot sizes differ (Sanders, 1989), when design is non-standard as in the Nelder wheel design (Parrott et al., 2012) or when spatial correlation is not expected to follow standard patterns due to external variation (Bakka et al., 2019).

In this study we focused on phenotypes that can be modelled with a Gaussian distribution only. However, the INLA R package enables seamless modelling of other distributions such as binomial, Poisson and others. Phenotypic observations that follow these types of distribtions arise in agriucultural field trials. For some models the only code change required is a switch of the family such as from `inla(…, family = “Gaussian”)` to `inla(…, family = “Poisson”)`. We refer the reader to Krainski et al. (2018) or Blangiardo and Cameletti (2015) for more details.

While the R package INLA enables flexible modelling of data from multiple trials and years this might usually require increasing the model complexity by accounting for trial specific residual variance or trial specific spatial parameters - by increasing the number of hyper-parameters. We have performed such an analysis with the real wheat data, where spatial variation in 2011 and 2012 trials differed substantially. While this can be accommodated with the R package INLA we highlight that the INLA method works best when the number of hyper-parameters is small, typically less than ten, and not exceeding 20. This limitation is due to the numerical integration of multi-dimensional posterior distribution of hyper-parameter in INLA (Rue et al., 2017). Since there is limited information to estimate hyper-parameters from a single trial, a parsimonious solution would be to group similar trials together and estimate hyper-parameters per group instead of per trial. This is what we did for the 2011 and 2012 trials with the real wheat data.

The main drawback with using R package INLA for analysing modern agricultural trials is that genome-wide marker data is highly-dimensional, which leads to dense systems of equations. INLA is based on numerical approximations and numerical methods for sparse matrices, and even though INLA can fit genomic models either via the genomic relationship matrix or via marker effects (Strandén and Garrick, 2009), there is substantial computational overhead to handle such models, which is not the case for the pedigree model which has a sparse precision matrix (Steinsland and Jensen, 2010; Henderson et al., 1984). This is why we chose to fit the genome-wide markers directly via the principal component approach, which is similar to the proposal of Ødegård et al. (2018). Another option would be to fit a model with individual genetic effects following VanRaden (2008), but with a genomic relationship matrix that uses dense-sparse partitioning into core and non-core individuals (Misztal, 2016). More research is required in this area to increase the usefulness of the R package INLA for the modern breeding applications.

Finally, since the INLA method implements Bayesian models, prior distributions had to be set for all parameters of the model. The marker variance estimates in the models for Chilean wheat data were quite small, and we expected this to be larger. Testing the same models using the penalized complexity priors (Simpson et al., 2017) increased the mean marker variance. However, we have used the default prior distributions in INLA for simplicity. It should be emphasized that using default priors is a choice as much as using any other prior or even using a specific distribution for the phenotype data. Setting a prior based on the knowledge about the process is likely to improve the inference. Choosing a prior distribution for parameters in the model is not always straightforward and more work is being done in the statistics community to improve this (Fuglstad et al., 2019).

## 5. Conclusion

This study showed how to fit established spatial models for analysing agricultural field trials using the open-source R package INLA. The results from the simulation study showed higher accuracy when spatial dependency was modelled and the highest increase in accuracy was reached using the discrete AR1⊗AR1 model and the continuous SPDE approach. Both models can be seamlessly fitted with the R package INLA, including joint modelling of multiple trials. The SPDE approach is suitable for agricultural field trials, especially for trials that cannot have a grid-like structure such as the Nelder wheel design used in tree breeding.

## Author contributions statement

MLS, GG and IS conceived and designed the analysis. MLS and GG contributed simulated data, and MLS performed the analysis. MLS wrote the manuscript, GG edited the manuscript, and IS and JH commented on the simulation design and the manuscript structure and content. All authors have read and approved the final manuscript.

## Compliance with ethical standards

The authors declare that they have no conflict of interest.

